# Allelopathy as an evolutionarily stable strategy

**DOI:** 10.1101/2021.08.04.455130

**Authors:** Rachel M. McCoy, Joshua R. Widhalm, Gordon G. McNickle

**Affiliations:** Purdue Center for Plant Biology, Purdue University, West Lafayette, IN 47907, USA; Department of Horticulture and Landscape Architecture, Purdue University, 625 Agriculture Mall Drive, West Lafayette, IN 47907, USA; Department of Botany and Plant Pathology, Purdue University, 915 West State Street, West Lafayette, IN 47907, USA

**Author notes:** Correspondence: G.G. McNickle, Center for Plant Biology and Department of Botany and Plant Pathology, Purdue University, 915 West State Street, West Lafayette, IN 47907, USA.

**Keywords:** allelopathy, game theory, evolutionarily stable strategy, modeling

## Abstract

In plants, most competition is resource competition, where one plant simply pre-empts the resources away from its neighbours. Interference competition, as the name implies, is a form of direct interference to prevent resource access. Interference competition is common among animals who can physically fight, but in plants, one of the main mechanisms of interference competition is Allelopathy. allelopathic plants release of cytotoxic chemicals into the environment which can increase their ability to compete with surrounding organisms for limited resources. The circumstances and conditions favoring the development and maintenance of allelochemicals, however, is not well understood. Particularly, it seems strange that, despite the obvious benefits of allelopathy, it seems to have only rarely evolved. To gain insight into the cost and benefit of allelopathy, we have developed a 2 × 2 matrix game to model the interaction between plants that produce allelochemicals and plants that do not. Production of an allelochemical introduces novel cost associated with synthesis and detoxifying a toxic chemical but may also convey a competitive advantage. A plant that does not produce an allelochemical will suffer the cost of encountering one. Our model predicts three cases in which the evolutionarily stable strategies are different. In the first, the non-allelopathic plant is a stronger competitor, and not producing allelochemicals is the evolutionarily stable strategy. In the second, the allelopathic plant is the better competitor and production of allelochemicals is the more beneficial strategy. In the last case, neither is the evolutionarily stable strategy. Instead, there are alternating stable states, depending on whether the allelopathic or non-allelopathic plant arrived first. The generated model reveals circumstances leading to the evolution of allelochemicals and sheds light on utilizing allelochemicals as part of weed management strategies. In particular, the wide region of alternative stable states in most parameterizations, combined with the fact that the absence of allelopathy is likely the ancestral state, provides an elegant answer to the question of why allelopathy rarely evolves despite its obvious benefits. Allelopathic plants can indeed outcompete non-allelopathic plants, but this benefit is simply not great enough to allow them to go to fixation and spread through the population. Thus, most populations would remain purely non-allelopathic.

## INTRODUCTION

Competition is ubiquitous in the natural world, as there are finite resources available in a given time and space^1–3^. Thus, competition generally reduces plant fitness when resources, such as light, space, water and nutrients are limiting^4,5^. This type of competition for finite resources is broadly named resource competition and occurs when organisms compete by simply reducing the availability of resources to other organisms^6^. Alternatively, interference competition occurs when one organism interferes with, and therefore reduces, the ability of the other to obtain a shared resource while not necessarily drawing down resource concentrations^6^. Animals routinely face interference competition as they can physically fight over resources^7^. Sessile plants primarily compete via resource competition. However, one of the major mechanisms of interference competition in plants is mediated chemically through allelopathy^8^. For example, one of the best documented examples is allelopathy by walnut trees (*Juglans* spp.), mediated by the allelochemical juglone, which is toxic to a variety of crop and horticultural species, including corn and soybean^12^ and tomato and cucumber^13^.

Allelopathy is the production of chemicals, called allelochemicals, that are released into the environment and negatively affect the growth and development of competing individuals^9^. Although the term was first used in 1937, the effect has been recognized for thousands of years^9^. Unfortunately, there have been difficulties in studying the competitive effects of allelopathy because of methodological difficulties. For example, for many years experiments used soil additives such as activated charcoal that were thought to prevent the activity of allelochemicals with the goal of comparing how plants grew either with or without the presence of this form of interference competition. Unfortunately, it was later learned that activated charcoal also stimulates nutrient availability, and thus, many years of research showing the negative effects of allelochemicals were probably just detecting the positive effects of fertilization (*e.g*.^10,11^).

Despite limitations in the ability to experimentally study allelopathy, it has been implicated in the success of some invasive plants, highlighting the advantage of interference competition as a strategy^14^. Invasion by non-native species is ranked the second strongest risk to natural diversity^15^. For example, Paterson’s curse (*Echium plantagineum* L.) is an invasive weed in Australia, affecting up to 30 Mha, whose invasion success is partially attributed to production of the allelochemical shikonin and its derivatives^16^. Indeed, one commonly invoked mechanism for invasion by non-native species is the novel weapons hypothesis, which suggests invasive species are successful through use of competitive strategies for which native species have not co-evolved counter strategies^17,18^. This mechanism has been linked to the invasion success of allelopathic Policeman’s helmet (*Impatiens glandulifera*)^19^, which releases a compound structurally similar to shikonin called 2-methoxy-1,4-naphthoquinone (2-MNQ) that elicits negative effects on herb germination and mycelium growth and is otherwise absent in soils without *I. glandulifera*, thus suggesting 2-MNQ may function as a “novel weapon”^19–21^. From these studies, it may be possible that allelochemicals may have significant potential for genetically modified cropping systems to enhance the competitive ability of crop species over weeds.

Despite the potential advantages of allelochemicals as an evolved tool for interference competition, they seem to have only rarely evolved. Here, we report an evolutionary game theoretic model to probe the benefits and circumstances that might favor the evolution of allelochemicals to better understand why they might not be more common in plants. Specifically, we ask: 1) What circumstances favor the production of allelochemicals? 2) How does the cost of producing an allelochemical affect fitness of the plant producing the allelochemical and plants competing with that plant? 3) When will allelopathic plants be stable in a population? Beyond the implications for evolutionary ecology, understanding the evolution of allelopathy has the potential to inform the design of applications for agriculture, from the integration of allelopathic crops into farming systems to the use of synthetic biology to create a crop that produces its own allelochemical-based weed control.

## MATERIALS AND METHODS

### Model development

We developed a 2 × 2 matrix game of interactions among a plant player with (*+A*) and without (−*A*) allelopathy. We assumed that competition creates benefits of available resources (B), that the cost (*C*) to the player of producing allelochemicals is the sum of the costs of production of the allelochemical and detoxification to prevent autotoxicity, and that allelochemicals impose some different cost to the opponent in the form of toxicity and/or detoxification (*T*). We further assumed that benefits were shared unequally, encompassed by a parameter, *a*, that represents the proportion of benefits the allelopathic plant receives when competing with a non-allelopathic plant. These parameters of the model should adhere to the following: 0 < *B*,*C*,*T* and 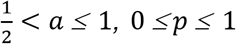, 0 *≤p ≤* 1 where *p* is the proportion of allelopathic plants in the population. We found this four-parameter model to be the simplest possible model that generates the evolution of allelopathy in ways that seem true to nature, though we describe two possible simpler alternatives in the Supplementary Information that explore how the parameters *a* and *T* individually shape model solutions. We understand the limitations imposed by the simplicity of the model, but the four, simple parameters encompass complex, multifaceted biological possibilities, and the simplicity allows us to ask large-scale questions about the ecology and evolution of allelopathy.

Combining these parameters, we can derive the payoff, *G*_*ν*,*u*_, across several competitive contexts where *ν* is the focal plant strategy (+*A* or −*A*), and *u* is the neighboring plant strategy (+*A* or −*A*). Finally, we also assume that there are two plants competing in something like a pot experiment, because we imagine this is the most likely way to empirically test our model in the future (e.g.^10,11^). However, the equations below can be extended to any number of competing plants by simply replacing 2 with *N*, where *N* is the number of competing plants.

First, when both plants produce allelochemicals, we argue that they will, on average, share the total benefit of the soil volume equally, 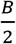, but will also pay the cost of producing and detoxifying allelochemicals, *C*. Thus, the fitness pay-off to a plant in a population of pure +*A* plants is:

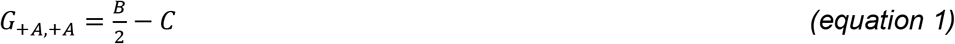

Second, in a mixed population of +*A* and −*A* plants, the +*A* plant will pay the cost *C* but will share the benefits *B* differently. Instead of equally sharing the benefits, the player will get a proportion of benefits, *a*, that takes into account the competitive advantage of production of allelochemicals according to:

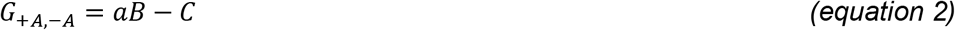

Inversely, in the mixed population, the −*A* plant obtains the remaining benefit, represented by (1−*a*)*B*, and pays the cost of toxicity, *T*, according to:

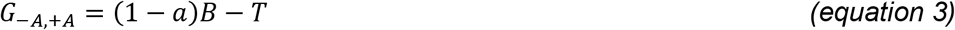

Finally, in a pure population of −*A* plants, because no plant produces allelochemicals, they merely share the benefits as 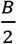 and have no costs associated with allelochemicals, according to:

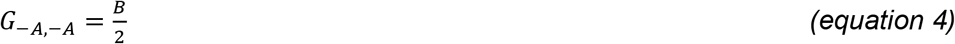

Combined, equations 1–4 yield the pay-off matrix shown as Figure 1.

**FIGURE 1:**
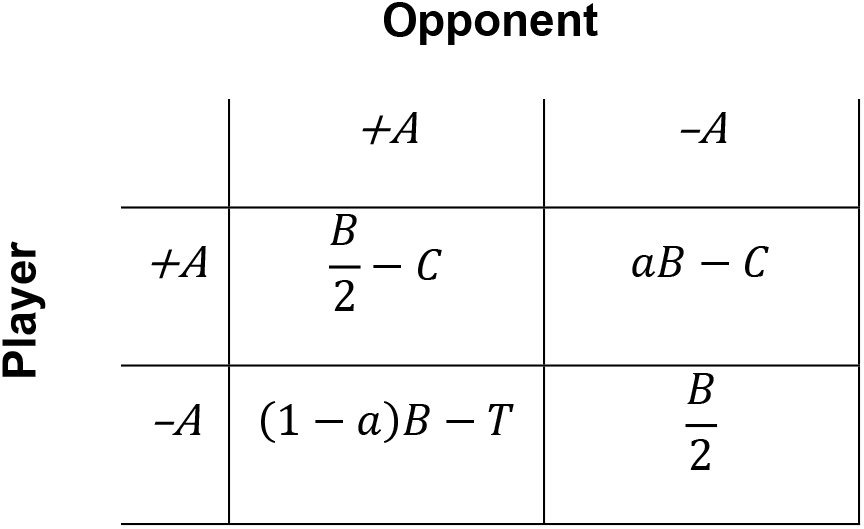
Symmetric pay-off matrix for competition between plants that either produce allelochemicals (+A) or not (-A). See text for parameter definitions.

### Evolutionarily stable strategy definition

In a matrix game, an evolutionarily stable strategy (ESS) is identical to a Nash equilibrium where a participant cannot gain by changing strategy if the other participant’s strategy does not change^22,23^. Thus, a pure ESS is defined as the strategy which once adopted by members of a population cannot be invaded by any alternative strategy. Mixed ESSs are also permissible where multiple strategies either ecologically coexist through evolutionary time or form non-coexisting alternative stable states (sometimes also called priority effects). Here, in a 2 × 2 matrix game, if *G*_*ν*,*u*_ is the fitness payoff of a focal plant species using strategy ‘*ν*’ against a competing plant species using strategy ‘*u*’ such that *ν* ≠ *u*, then *ν* is a pure ESS if and only if: *G*_*ν*,*ν*_ > *G*_*u*,*ν*_ and *G*_*ν*,*u*_ > *G*_*u*,*u*_. Alternatively, *u* is a pure ESS when *G*_*u*,*ν*_ > *G*_*ν*,*ν*_ and *G*_*u*,*u*_ > *G*_*ν*,*u*_ (*i.e*. under the opposite inequalities). Most interestingly, under this definition mixed ESS solutions are possible where the two strategies may coexist or for a system of alternative stable states ^24^. A mixed ESS occurs when: *G*_*u*,*ν*_ > *G*_*ν*,*ν*_ and *G*_*ν*,*u*_ > *G*_*u*,*u*_. Alternative stable states occur when: *G*_*ν*,*ν*_ > *G*_*u*,*ν*_ and *G*_*u*,*u*_ > *G*_*ν*,*u*_. Together, a keen observer will note that these four inequalities form all possible pairs of inequalities within each column of a 2 × 2 payoff matrix as drawn here (Figure 1).

## RESULTS

### Pure evolutionarily stable strategies

For +*A* to be a pure ESS, +*A* needs to be able to (i) invade a population of −*A* and (ii) resist invasion from −*A*. According to the ESS definition, this occurs when:

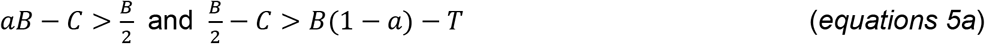

Equations 5a can be rearranged into isoclines in *B* and *C* space to find:

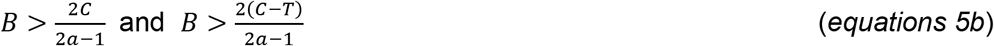

Alternatively, for −*A* to be ESS, −*A* needs to be able to (i) invade a population of +*A* and (ii) resist invasion from +*A*.

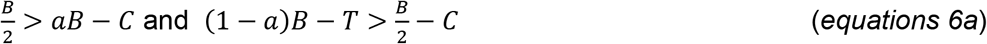

Equations 6a can be rearranged to find:

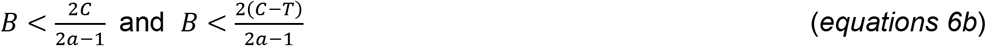

Notice that equations 5 and 6 are simply opposite inequalities.

### Mixed evolutionarily stable strategies

The mixed strategy, where there are alternating stable states such that either −*A* or +*A* can resist invasion from the other strategy occurs when,

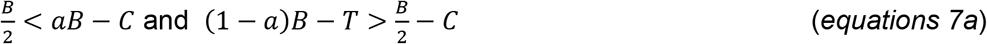

Equations 7a can be rearranged to find:

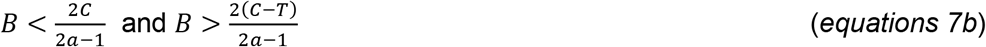

The isoclines in equations 5-7 create two parallel lines, each with slope 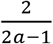 but that either intercept the y-axis at 0 or at 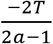. Thus, depending on the values of *a* and *T*, we can plot the entire solution space graphically in positive *B* and *C* phase space (Figure 2). For +*A* to be the ESS, the parameters need to be above both isoclines. For −*A* to be the ESS, the parameters need to be below both isoclines. Between the two lines, which will never cross as they have the same slope, there is a region of alternative stable states, also sometimes called a priority effect. In the region of alternative stable states, either strategy might occur, but the answer depends on the history of the system. That is, whichever strategy was there first becomes the ESS>

**FIGURE 2:**
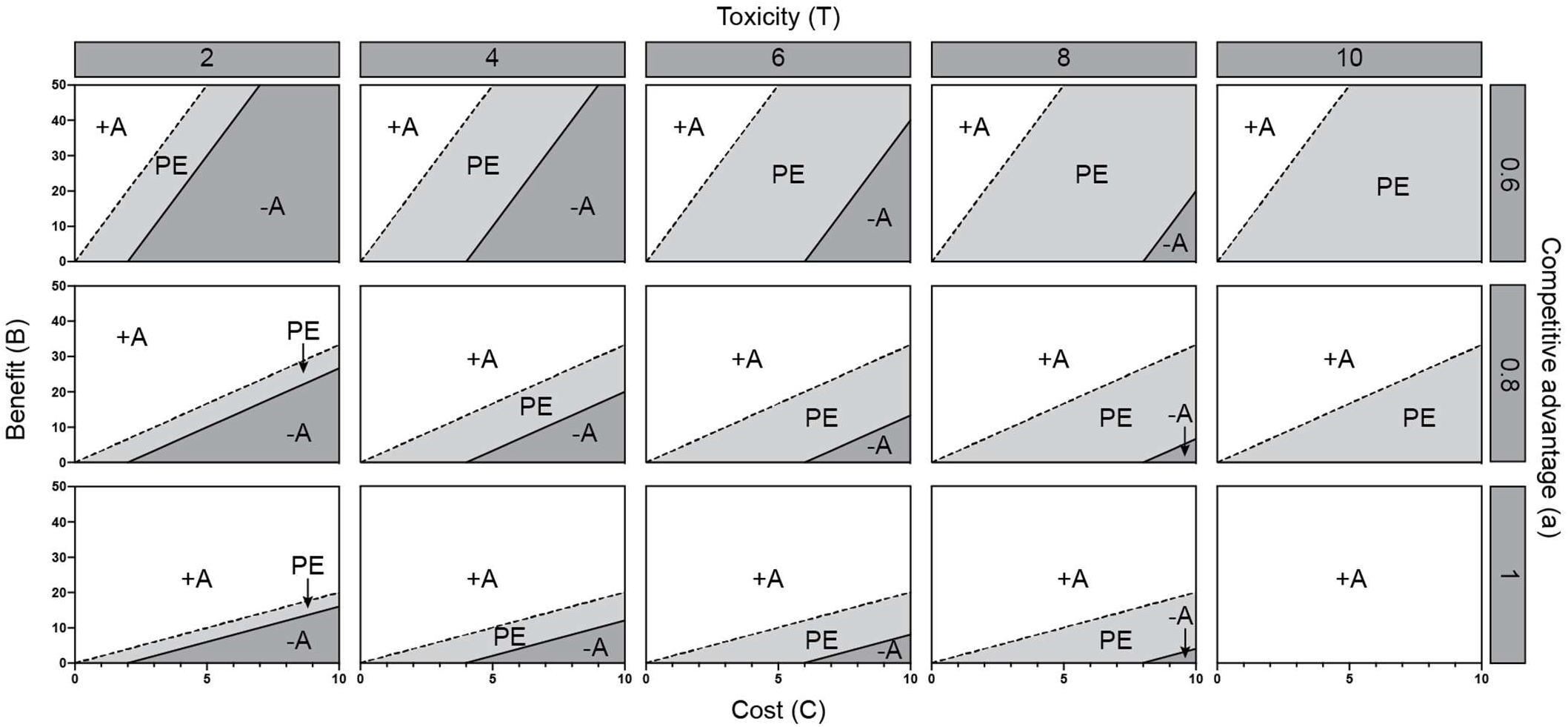
Isoclines that depict ESS states in B and C phase space depending on the value of T 2C (columns) or a (rows). The dashed line represents the isocline 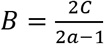. The solid line represents the isocline 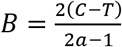. White space is the area of parameter space where production of allelochemicals (+A) is the ESS. Dark grey is where not producing allelochemicals is the ESS (-A). The space in between (light grey) is where priority effect (PE) occurs.

In our model, coexistence is never possible. For it to be so, would require:

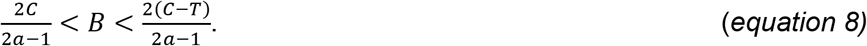

Because *T* > 0 by definition, these conditions can never be met. This suggests that within a population, all plants of a species will either produce or not produce allelochemicals.

Allelopathic plants gain a competitive advantage only when 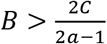 which can offset the cost of producing allelochemicals beyond just the cost of toxicity on the neighboring plant (Figure 2). However, if 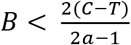 then non-allelopathic −*A* plants gain the competitive advantage because the benefits of allelopathy do not outweigh the costs to the allelopathic plant, or the allelopathic chemical is simply not toxic enough to generate a benefit (*i.e*. low *T*). This region of pure +*A* as the ESS expands as *a* increases (Figure 2). When *a* > 0.5, but 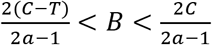, there is an interesting region between the two isoclines of alternative stable states where only one strategy can exist at a time, but which one occurs depends on the initial conditions (*i.e*. on this history of colonization and/or mutation). This area of alternative stable states also expands with increasing *T* but decreasing *a*.

## DISCUSSION

In this study, we developed and analyzed a model of the evolution of allelopathy between two competing plants as an evolutionary game to examine the conditions under which the production and deployment of allelochemicals becomes a favorable competitive strategy. The model has four simple parameters that describe costs and benefits among players. Somewhat intuitively, allelopathy can only evolve when the benefits to the allelopathic plant outweigh the costs, but the model outlines these precise conditions in 4-dimensional phase space (Figure 2). For example, in the case of an extremely toxic allelochemical (*i.e*. large *T*) that also happens to be metabolically costly to produce (*i.e*. large *C*), the model makes it clear that there must be relatively high benefits (e.g. high *B*, very fertile environments) and confer a very large competitive advantage (large *a*). Indeed, except where *a* approaches 1 and *T* is large, we see large regions of alternative stable states, and relatively small regions where +*A* is the pure ESS. Assuming that −*A* is the ancestral condition, we argue that this might explain why allelopathy has been relatively rare to evolve, despite the obvious advantage. That is, in the region of alternative stable states, any +*A* mutants would simply not be able to invade the ancestral −*A* population because of their priority effect advantage. The relative rarity of allelopathy in nature might indicate natural environments found on this planet exist closer to the upper left region of Figure 2, though future work should investigate whether the biochemical cost of production and the cost of detoxification are substantial energetic costs to plants to narrow down the region of parameter space that exists in natural plant communities. There may be some biochemical constraints that place the plant kingdom in this part of the phase space, and this would be an important area for plant biologists to explore further. It is also possible that allelopathy is more common than current knowledge suggests.

One way of reducing the cost of producing novel allelochemicals, *C*, is to harness existing metabolic frameworks. Many species producing naphthoquinone-based compounds, for example, have independently evolved to do so from 1,4-dihydroxy-2-naphthoic acid (DHNA), an intermediate of the phylloquinone (vitamin K1) pathway^32,33^. Examples include juglone in black walnut trees^34^, lawsone and 2-MNQ in the Balsaminaceae (e.g. *Impatiens* species)^35^, lawsone and lapachol in the Bignoniaceae^36^, anthraquinones like alizarin made by Rubiaceae species^37^, and anthrasesamones produced by sesame (*Sesamum indicum*, Pedaliaceae)^38^. Interestingly, juglone, lawsone, and 2-MNQ are all implicated as allelochemicals^19,39,40^. This indicates that DHNA derived from the phylloquinone pathway, which is present in all plants, likely provides a lower cost path for plants to synthesize allelochemicals.

Over time, the cost of allelochemical toxicity, *T*, to −*A* plants could be mitigated by evolution of mechanisms to tolerate or detoxify the allelochemical. Therefore, the competitive disadvantage of not producing the allelochemical to −*A* plants would dissipate; however, the cost of detoxification, which is also part of *T*, would likely remain non-zero. We hypothesize that the evolution of *T* can draw inferences from evolution of herbicide resistances in plants, which occur via mutations in herbicide target sites (target-site resistance) or non-target sites (non-target-site resistance)^41^. In an analogous scenario of non-target-site resistance, the allelochemical itself or the toxicity arising from the allelochemical could be metabolically counteracted through biochemical modification and/or compartmentalization of the allelochemical or its modified product. Thus, non-target-site resistance is referred to as “metabolism-based resistance”^42^. Metabolism-based resistance to herbicides is primarily achieved via four gene families: cytochrome P450 monooxygenases, glutathione transferases (GSTs), glycosyltransferases, and/or ABC transporters^43^. It is likely that plants that evolve in proximity to an allelopathic plant use similar methods to tolerate allelochemicals. GSTs function by covalently linking glutathione (GSH) with compounds that are hydrophobic and electrophilic^44^; some also function as carriers that transport GSH-conjugates to vacuoles for detoxification^45^. Black-grass (*Alopercurus myosuroides*) is a weed species that has evolved resistance to multiple herbicides by over expressing a single GST, AmGSTF1. Heterologous overexpression AmGSTF1 in *Arabidopsis thaliana* was shown to be sufficient to confer resistance to multiple herbicides^46^. Moreover, Arabidopsis seedlings grown *in vitro* in the presence of GSH in juglone-containing media were found to display root growth phenotypes indistinguishable from wild type (Meyer et al 2020). Beyond conjugation with GSH, glycosylation appears to be a major mechanism of detoxification of specialized metabolites^47^. Indeed, much of the juglone found in black walnut is glycosylated^48^, suggesting that one of the mechanisms black walnut uses to tolerate producing and storing an autotoxic compound is through glycosylation. Reduced uptake or increased export could also confer some tolerance to allelopathic exposure. Mutations in transport proteins have been shown to confer resistance to herbicides through decreased uptake (reviewed in^49^). Additionally, fungi and bacteria have been shown to be able to degrade structurally diverse, toxic chemicals from a variety of plant families^50–52^. Studies from microorganisms may provide more insight into mechanisms plants use to tolerate allelochemicals or provide guidance for transgenic strategies to convey resistance to allelochemicals.

### Model assumptions and caveats

As ever, any model comes with some caveats. One big one, is that factors which are not included in our model can affect Allelopathy. For example, in natural environments, allelopathy is affected by the ecology of the soil^25^. *Pseudomonas* J1, a soil bacteria isolated from soil surrounding a black walnut, is capable of growing on juglone as its sole carbon source^26^. Further, ailanthanone from *Ailanthus altissima* is more effective at suppressing growth of radish in sterile soil^27^. These and other studies show that degradation of allelochemicals by soil microbes is a factor in the toxicity of an allelochemical in a given environment. Such degradation would lead to a decrease in *T*, indicating that the parameter *T* for the same compound could be different in different soils and ecosystems. Similarly, the soil microbiome has been shown to have diverse effects on plant fitness (reviewed in^28^). In addition to directly harming nearby plants, allelopathy may also play a role in altering the microbial soil community to the benefit of the allelopathic plant.

Another example of a factor absent from our model is how Allelopathy interacts with other plant interactions. For example, invasive garlic mustard has been shown to inhibit the interaction between seedlings of competitors and their mutualistic fungi^29^. Similarly, *I. glandulifera* invasion disrupts symbiotic associations between arbuscular mycorrhiza and native saplings^20^, likely via the release of 2-MNQ, which was also shown to inhibit mycelium growth of ectomycorrhiza fungi^19^. Conversely, some studies have suggested that the plant microbiome reduces the effect of allelochemicals on the plant^30^, in effect lowering the cost of toxicity/detoxification, *T*, to opponents. *Paxillus involutus*, a mycorrhizal fungi of black spruce (*Picea mariana*), has been shown to be able to degrade allelopathic compounds produced by *Kalmia angustifolia*, perhaps conveying some tolerance to black spruce^31^. These examples demonstrate the complexity of studying allelopathy in field conditions that our model does not capture.

### Implications and applications of the model

Investigating the means by which plants reduce cost and increase fitness in the presence of allelochemicals will allow more predictable integration of allelopathy as part of weed management strategies in cropping systems. For example, intercropping is a common agricultural practice used in many parts of the world to improve land use efficiency, to mitigate the risk of a single crop failing, and to diversify farming income. Often, intercropping involves co-cultivation of two or more cash crops, but in some cases a cash crop is grown alongside a non-cash crop to provide benefits, such as weed suppression, to the primary crop^53^. In either case, intercropped species are grown in close enough proximity to allow biological interaction. Therefore, the allelopathic potential of each species should be considered when designing mixed cropping systems^54^. For example, a study by Iqbal *et al.^55^* showed that intercropping cotton (*Gossypium hirsutum* L., cv FH901) with allelopathic crops, including sorghum (*Sorghum bicolor* L.), soybean (*Glycine max* L.), or sesame (*Sesamum indicum* L.), was an effective strategy to control purple nutsedge (*Cyperus rotundus* L.), a common aggressive weed found in parts of South Asia. According to the matrix game presented here, the fitness pay-off to both cotton and purple nutsedge (the −*A* species) would be expected to decrease as the toxicity, *T*, of allelochemicals produced by sorghum, soybean, or sesame (the +*A* species) increased. Indeed, seed cotton yield was found to decrease between 8-23% in all intercropping systems, compared to unmanaged cotton alone. Similarly, the presence of allelopathic species led to 70-96% reduced purple nutsedge density^55^. That control of purple nutsedge was found to be more effective in the second year of the study compared to the first year, which was suggested to be the result of residual allelochemicals leftover in the soil in year two^55^. This is consistent with purple nutsedge paying an increased penalty, *T*, to detoxify higher levels of allelochemicals.

Herbicide applications have increased over the last 25 years in many major cropping systems^56^. With this trend, so too has the number of weeds that have developed resistance to commonly used pesticides^57^. To address the lack of new herbicidal modes of action needed to combat resistant weeds^58^, allelochemicals, which offer a wide diversity of new chemical structures, have been suggested as sources for developing novel herbicides^59^. One attractive strategy is to engineer or breed production of allelochemicals into non-allelopathic cash crops, although autotoxicity and the metabolic cost of biosynthesis must remain low enough to not significantly impact agricultural performance^60^. If the cost, *C*, to the crop engineered to be +*A* is too high then it would not be an ESS and could be invaded by-A species (*i.e*. weeds) (equations 5a and 5b). At the same time, if the cost, *C*, to the +*A* crop is too low then it may allow it to become too easily capable of escaping and invading native populations of −*A* species (Figure 2). If the crop were engineered to produce and detoxify an allelochemical such that it was in the realm of alternative stable states, the allelopathic crop would be able to resist invasion from −*A* weeds, without the possibilty of escape and invasion. Conversely, by purposefully engineering a less fit crop to fall *outside* the region where +*A* is ESS (Figure 2), it would also provide a mechanism by which to prevent escape of the transgenic species. Such application could be useful in cover cropping where certain cover crops that not controlled prior to planting cash crops can become weeds.

Finally, another interesting consideration in the evolution of allelopathy is the presence of allelobiosis. Allelobiosis is a relatively new term that describes communication between plants via non-toxic compounds^61^. For example, planting tomato (*Lycopersicom esculentum*) in proximity with sagebrush (*Artemisia tridentata*) resulted in increased production of proteinase inhibitors in tomato due to methyl jasmonate released by the sagebrush^62^. Though allelobiosis has been most often demonstrated with volatile compounds, there are examples of this kind of plant-plant communication through the rhizosphere. Indeed, Li *et al*. (2016) found that allelobiosis and allelopathy coexist in interactions of weeds with allelopathic wheat. Root exudates from weed species were sufficient to induce allelopathy in the wheat, suggesting a chemical signal sensed by the wheat. Further work is necessary to detangle the effects of allelobiosis and allelopathy, especially in the case of inducible production of allelochemicals. As more information arises, alelobiosis could be an important factor to include in future efforts to expand the modeling of Allelopathy as an ESS>

### Conclusion

Our model predicts three ESS cases, differing in the benefit and cost to the allelopathic plant. In the first, the non-allelopathic plant is a stronger competitor due to high metabolic costs to the allelopathic plant, and not producing allelochemicals is the evolutionarily stable strategy. In the second, the allelopathic plant is the better competitor and production of allelochemicals is the more beneficial strategy. In the last case, the allelopathic and non-allelopathic plants are equal competitors, but pay different costs resulting in alternative stable states depending on the history of the system. We find that despite the obvious benefits of allelopathy, there are relatively few conditions that lead to +*A* as a pure ESS, and that if −*A* is the ancestral state the large regions dominated by priority effects would mean +*A* mutants cannot successfully spread in a population. We argue that these results potentially help explain the relative rarity of allelopathy in nature. Additionally, the four parameters give insight into molecular mechanisms that future biochemical and molecular work could seek to better understand. Further empirical exploration of this model could lead to useful agricultural tools.

## AUTHORS’ CONTRIBUTIONS

R.M.M. conceived the project with guidance from G.G.M. and J.R.W.; R.M.M. and G.G.M. performed the modeling; R.M.M., J.R.W., and G.G.M. analyzed the model solutions and wrote the paper.

## ACKNOWLEDGEMENTS

This work was supported by the USDA National Institute of Food and Agriculture Predoctoral Grant 2018-67011-28032 to R.M.M. and start-up funds from Purdue University to G.G.M. and J.R.W. This work was also supported by the USDA National Institute of Food and Agriculture Hatch Projects number 1010722 to G.G.M. and number 177845 to J.R.W.

## COMPETING INTERESTS

The authors declare no competing interests.

## SUPPLEMENTARY INFORMATION

The main text describes a four parameter matrix game of Allelopathy. This model includes two different features of the allelochemical: (i) how it increases fitness benefits to the allelopathic plant through increased competitive ability (the parameter *a*), and; (ii) how it imposes costs on competitors through toxicity (the parameter *T*). Here, we examine two additional three parameter models, one without *a*, and one without *T*, to further probe whether this two-feature way of describing allelochemicals was necessary.

### Exclusion of the parameter *a*

In the main text, the parameter *a* signifies the competitive advantage conveyed to allelopathic plants when competing for resources. Here, we examine a three-parameter model without the inclusion of the parameter *a* which we began with. The solution to this game showed us that this three parameter model was too simple, and we therefore developed the four parameter model described in the main text. In this version, we assume that competing plants share the benefit in a given area equally (*i.e*. 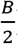). This yields the following pay-off matrix:

**Supplementary Figure S1:**
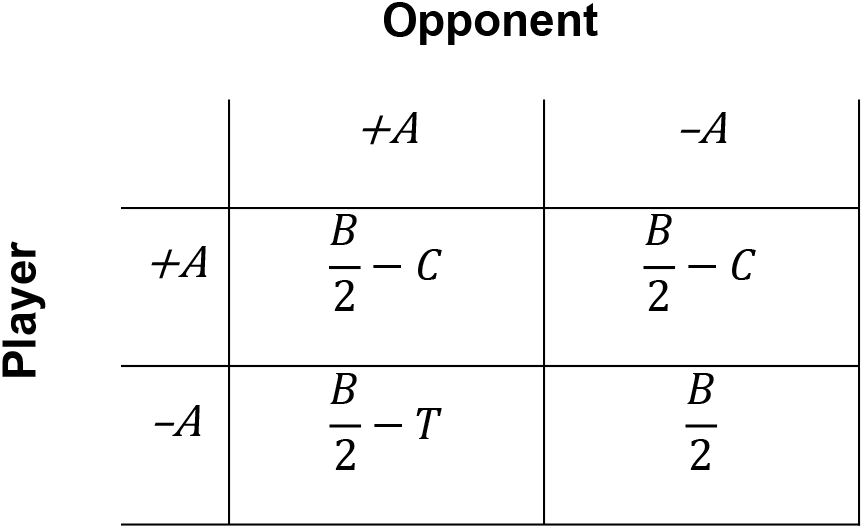
Symmetric pay-off matrix for competition between plants that either produce allelochemicals (+A) or not (-A) without the inclusion of the competitive parameter a

In this case, for +*A* to be a pure ESS, (i) +*A* needs to be able to invade a population of −*A* and (ii) needs to resist invasion from −*A*. According to the ESS definition, this occurs when:

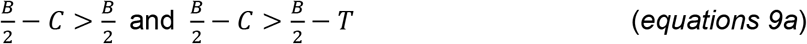

Both of which can be rearranged to:

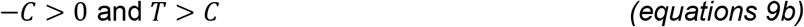

Therefore, +*A* can never be the ESS in this simpler three parameter version of the game because *C* is by definition greater than zero, so the first condition in equations 9b cannot be met.

Conversely, for −*A* to be a pure ESS, (i) −*A* needs to be able to invade a population of +*A* and (ii) needs to resist invasion from +*A*. According to the ESS definition, this occurs when:

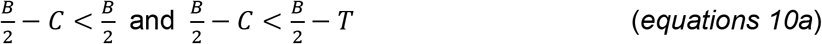

Both of which can be rearranged to:

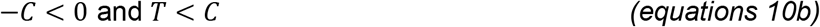

In this case, *B* has no bearing on the ESS. The only factors that matter are the relative values of *C* and *T*. Also, allelopathy can never be the ESS, which seems counterintuitive, and incorrect since we find allelopathic plants in nature. Thus, it seems clear that if the only thing an allelochemical does is impose a fitness cost on competitors, this is not sufficient for alleopathy to evolve.

### Exclusion of the parameter *T*

In the main text the parameter *T* signifies the toxicity of the allelochemical. Here, we examine a simpler three-parameter model without the inclusion of the parameter *T*, the pay-off matrix is as follows:

**Supplementary Figure S2:**
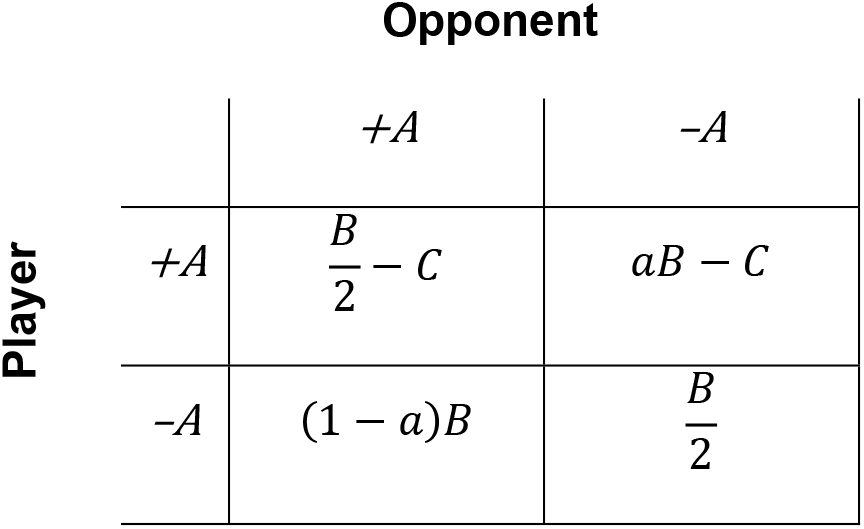
Symmetric pay-off matrix for competition between plants that either produce allelochemicals (+A) or not −A) without the inclusion of T.

In this case, for +*A* to be a pure ESS, (i) +*A* needs to be able to invade a population of −*A* and (ii) needs to resist invasion from −*A*. According to the ESS definition, this occurs when:

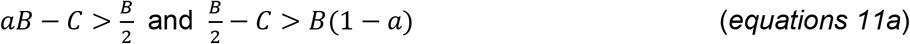

Both of which can be rearranged to:

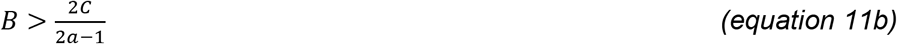

Conversely, for −*A* to be a pure ESS, (i) −*A* needs to be able to invade a population of +*A* and (ii) needs to resist invasion from +*A*. According to the ESS definition, this occurs when:

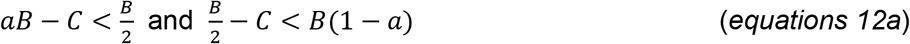

Both of which can be rearranged to:

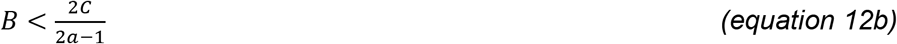

Because there is only a single isocline (equations 13b and 14b), mixed ESS are not possible. In this situation, only pure ESS solutions can exist. Above the isocline 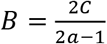, +*A* is ESS and below −*A* is ESS (Supplementary figure S3).

**Supplementary Figure S3:**
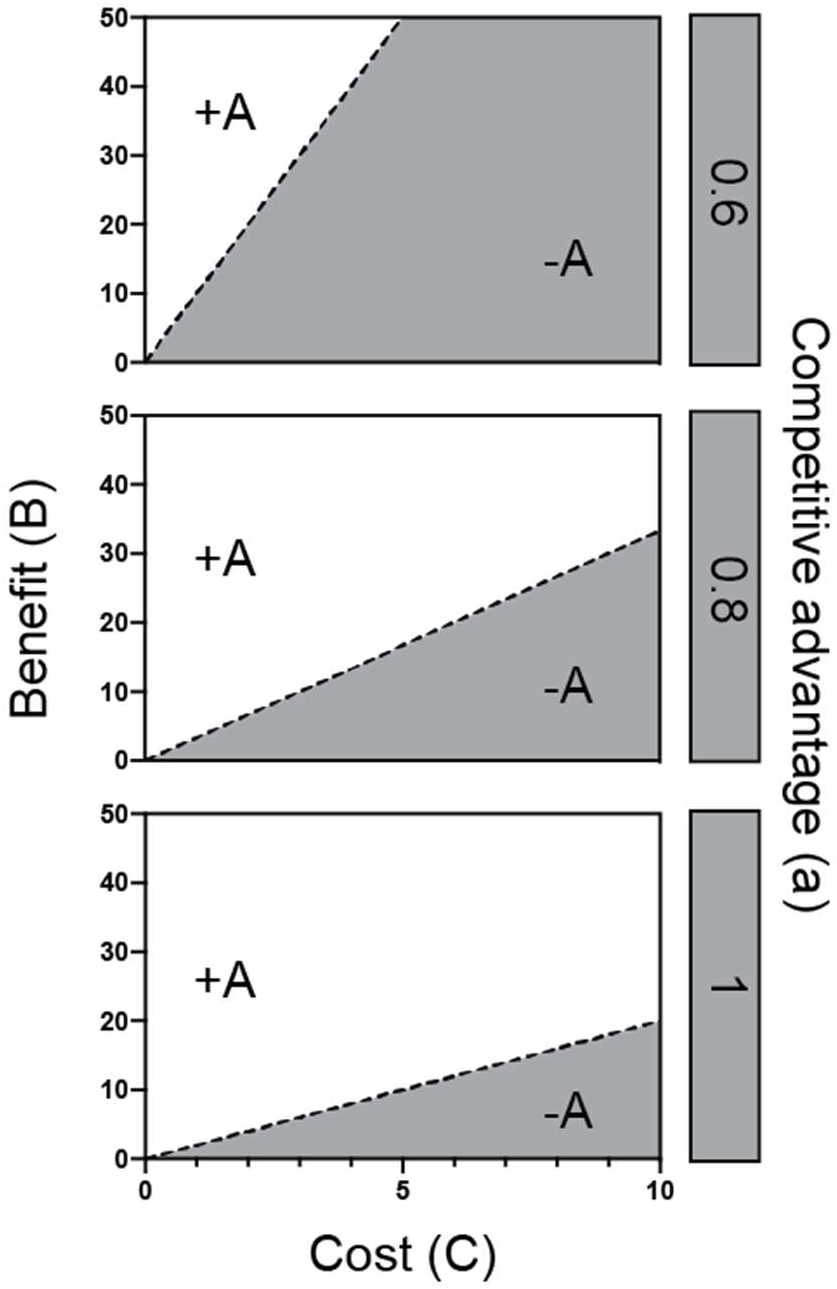
Isoclines that depict ESS states in B and C phase space depending on the value a (rows). The dashed line represents the isocline 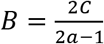. White space is the area of parameter space where production of allelochemicals (+A) is the ESS. Dark grey is where not producing allelochemicals is the ESS (-A).

Interestingly, a model that only includes a fitness benefit to the focal plant that emerges from alleopathy could be a useful model of allopathy. It lacks the priority effects predicted by the four parameter model in the main text (Fig 2), which presents a testable hypothesis. However, given that the main biological feature of alleopathy is the toxicity that they cause to neighbours, we opted to include *T* in the model described in the main text, even though this model shows that this toxicity is not strictly necessary.

## Notes

### Competing Interest Statement

The authors have declared no competing interest.

